# Host-symbiont population genomics provide insights into partner fidelity, transmission mode and habitat adaptation in deep-sea hydrothermal vent snails

**DOI:** 10.1101/2021.07.13.452231

**Authors:** Corinna Breusing, Maximilian Genetti, Shelbi L. Russell, Russell B. Corbett-Detig, Roxanne A. Beinart

**Author notes:** Corinna Breusing, Roxanne A. Beinart **Email**. **Author Contributions**: C.B. and R.A.B. designed the study with advice from S.L.R. and R.C.D. C.B. performed data analyses and wrote the first draft of the paper. M.G. prepared Illumina sequencing libraries. All authors contributed to data interpretation and writing of the manuscript. **Competing Interest Statement**: The authors declare no competing interests. **Classification**: Biological Sciences, Evolution.

## Abstract

Symbiont specificity, both at the phylotype and strain level, can have profound consequences for host ecology and evolution. However, except for insights from a few model symbiosis systems, the degree of partner fidelity and the influence of host versus environmental factors on symbiont composition are still poorly understood. Nutritional symbioses between invertebrate animals and chemosynthetic bacteria at deep-sea hydrothermal vents are examples of relatively selective associations, where hosts affiliate only with particular, environmentally acquired phylotypes of gammaproteobacterial or campylobacterial symbionts. In hydrothermal vent snails of the sister genera *Alviniconcha* and *Ifremeria* this phylotype specificity has been shown to play a role in habitat distribution and partitioning among different holobiont species. However, it is currently unknown if fidelity goes beyond species level associations that might influence genetic structuring, connectivity and habitat adaptation of holobiont populations. We used metagenomic analyses to assess sequence variation in hosts and symbionts and identify correlations with geographic and environmental factors. Our analyses indicate that host populations are not differentiated across a ~800 km gradient, while symbiont populations are clearly structured between vent locations due to a combination of neutral and selective processes. Overall, these results suggest that host individuals flexibly associate with locally adapted strains of their specific symbiont phylotypes, which supports a long-standing but untested paradigm of the benefits of horizontal transmission. Strain flexibility in these snails likely enables host populations to exploit a range of habitat conditions, which might favor wide-spread genetic connectivity and ecological resilience unless physical dispersal barriers are present.

**Significance Statement:** Symbiont composition in horizontally transmitted symbioses is influenced by a combination of host genetics, environmental conditions and geographic barriers. Yet the relative importance of these factors and the effects of adaptive versus neutral evolutionary forces on symbiont population structure remain unknown in the majority of marine symbioses. To address these questions, we applied population genomic approaches in four species of deep-sea hydrothermal vent snails that live in obligate association with chemosynthetic bacteria. Our analyses show that host genetics plays a minor role compared to environment for symbiont strain composition despite specificity to symbiont species and corroborate a long-standing hypothesis that vent invertebrates affiliate with locally adapted symbiont strains to cope with the variable habitat conditions characterizing hydrothermal vents.

## Introduction

Mutualistic relationships between eukaryotes and bacterial microbes are ubiquitous in nature. Symbionts enable hosts to gain access to novel resources and habitats, provide protection against pathogens and predators, and can be essential for the host’s diet (1, 2). For symbiotic associations to persist over evolutionary time, hosts must successfully transmit their symbionts from one generation to the other, either through symbiont acquisition from the environment (horizontal transmission), direct inheritance of symbiont lineages through the host germline (vertical transmission) or a combination of both mechanisms (mixed transmission) (3, 4). The mode of transmission has significant implications for the composition and variation of symbionts within and between host individuals. Vertical transmission typically results in strong genetic coupling between host and symbiont lineages and an accompanied reduction in intra-host symbiont diversity (3). By contrast, horizontal transmission exposes aposymbiotic hosts to a potentially heterogenous environmental pool of symbiont lineages (3), which can promote the formation of generalist partnerships, where multiple hosts and symbionts associate with each other, to more specialized associations between only one or a few potential partners (1, 5). This range is often referred to as host-symbiont specificity, which can vary in its taxonomic level for both partners depending on the symbiotic system. The degree of partner fidelity and its effect on symbiont composition can have dramatic impacts on holobiont functioning (6). For example, host-symbiont specificity, both at the species and genotype level, is crucial for light production in bioluminescent squid (7), while shifts in microbiome assemblages have been shown to affect phytoplankton growth rates (8) and the efficiency of nitrogen fixation in nodule-forming legumes (9–11).

Environmental transmission of obligate symbionts is particularly common in marine ecosystems (2). However, despite its importance for host biology, the relative contributions of host genetic, environmental and geographical factors to symbiont composition remain understudied in most horizontally transmitted marine symbioses. It has long been hypothesized that horizontal transmission in marine symbioses enables host organisms to associate with locally adapted symbiont strains, conferring fitness advantages in spatially and temporally variable marine habitats (3, 12–14). Especially for long-dispersing aposymbiotic larvae that are likely to encounter new habitat conditions when they settle, association with a locally adapted symbiont strain may be advantageous compared to carrying a vertically transmitted symbiont that might be maladapted at a non-native site. This hypothesis has been indirectly supported by evidence that marine animals with horizontally transmitted obligate microbial symbionts often host location-specific strains (14–19). However, the influence of local adaptation relative to neutral evolutionary processes on symbiont geographic structure and genomic traits has not been formally evaluated.

Chemosynthetic animal-microbe symbioses are globally significant phenomena that dominate hydrothermal vent and hydrocarbon seep ecosystems in the deep sea. In these associations, the bacterial partner uses chemical energy from the oxidation of reduced fluid compounds, such as hydrogen, sulfide or methane, to synthesize organic matter, which serves as primary nutrition for the host (20). Vent animals harboring chemosynthetic symbionts are typically highly selective in their partner choice: In the predominant number of cases host individuals associate with only 1–2 phylotypes (species or genera) of gammaproteobacterial or campylobacterial symbionts (20), whereas symbionts can exhibit a comparatively broad host range. Chemosynthetic endosymbioses in deep-sea snails of the Indo-Pacific sister genera *Alviniconcha* and *Ifremeria* are examples of reciprocally relatively specific partnerships, where host species harbor only particular symbiont species or genera across their geographic distribution. Given the absence of host-symbiont phylogenetic concordance, the symbionts are assumed to be environmentally acquired (21), although a pseudo-vertical transmission component is possible in *Ifremeria* given its brooding reproductive mode (22). In the Eastern Lau Spreading Center (ELSC), previous work suggested that specificity to functionally distinct symbiont phylotypes drives local and regional-scale habitat partitioning among four co-occurring *Alviniconcha* and *Ifremeria* species (23–26). *Alviniconcha boucheti* from the ELSC contains a campylobacterial phylotype (Epsilon) and is usually found at northern vent sites with high concentrations of sulfide and hydrogen, while *A. kojimai* and *A. strummeri* associate with different gammaproteobacterial phylotypes (Gamma1 and Gamma1/GammaLau, respectively) and usually occupy mid-latitude to southern vent sites where the concentrations of these chemical reductants are lower (23, 25–27). *Ifremeria nautilei* establishes dual symbioses with thiotrophic and methanotrophic gammaproteobacterial endosymbionts (27) and is co-distributed with *Alviniconcha* across their geographical range, although it typically segregates into habitat patches with reduced fluid flow relative to its sister genus (28). While it has been well-established that niche differentiation across hydrothermal vents is likely mediated by symbiont phylotype specificity in *Alviniconcha* and *Ifremeria* host species, nothing is known about the fidelity of these associations at the population level, how strain-level specificity might influence host population structure and connectivity, and how regional adaptation is conferred functionally.

In this study we applied population genomic methods to assess symbiont strain-level genetic variation and patterns of host-symbiont genetic subdivision in *Alviniconcha* and *Ifremeria* species from the ELSC and Tonga Volcanic Arc (Fig. 1; Table 1). Using multivariate statistical analyses, we evaluated the impact of host traits, environment and geography on symbiont composition in populations of both genera and assessed the effect of local adaptation on symbiont geographic structure.

**Figure 1.**
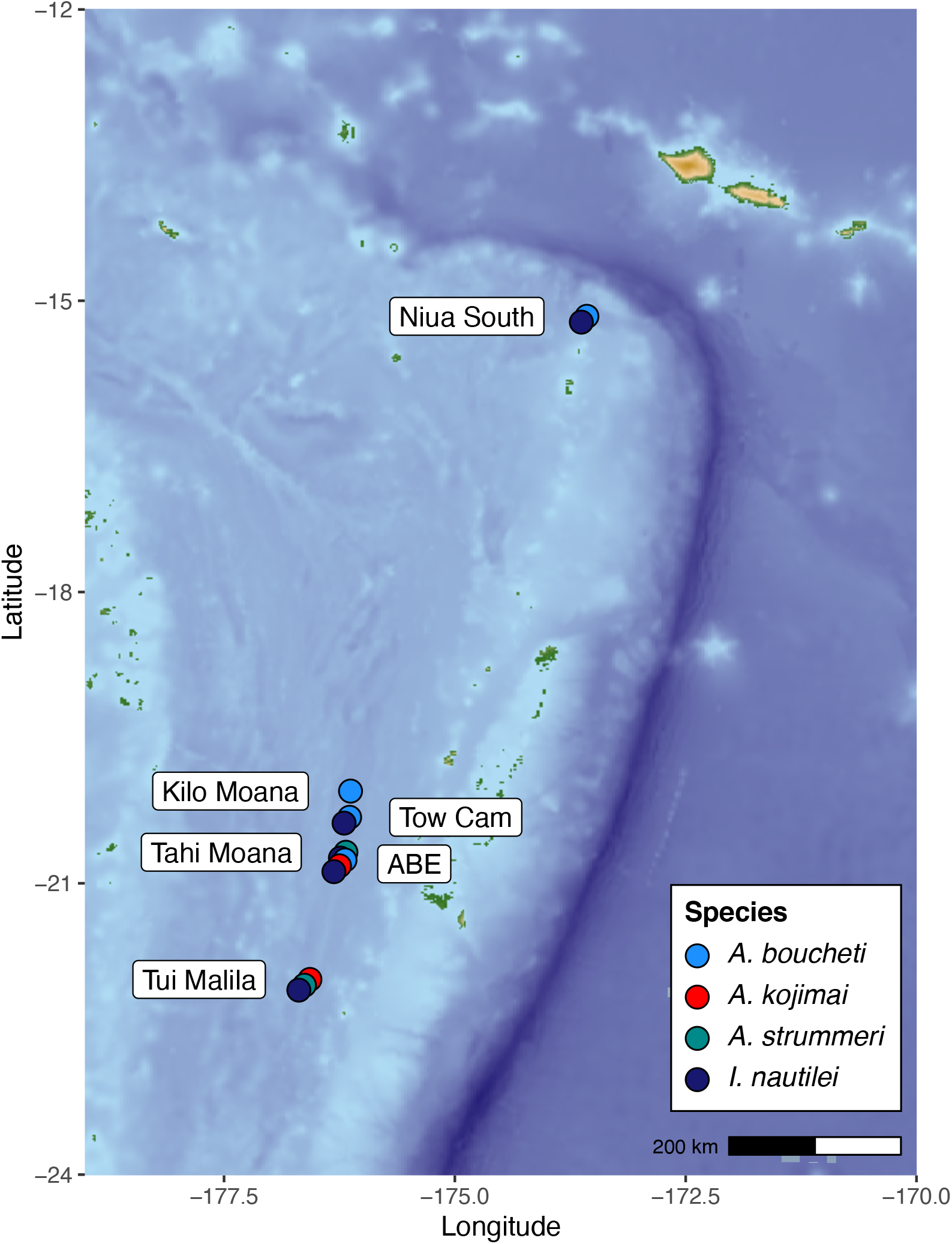
Sampling map for *Alviniconcha* and *Ifremeria* species in the Eastern Lau Spreading Center and Tonga Volcanic Arc.

**Table 1.**
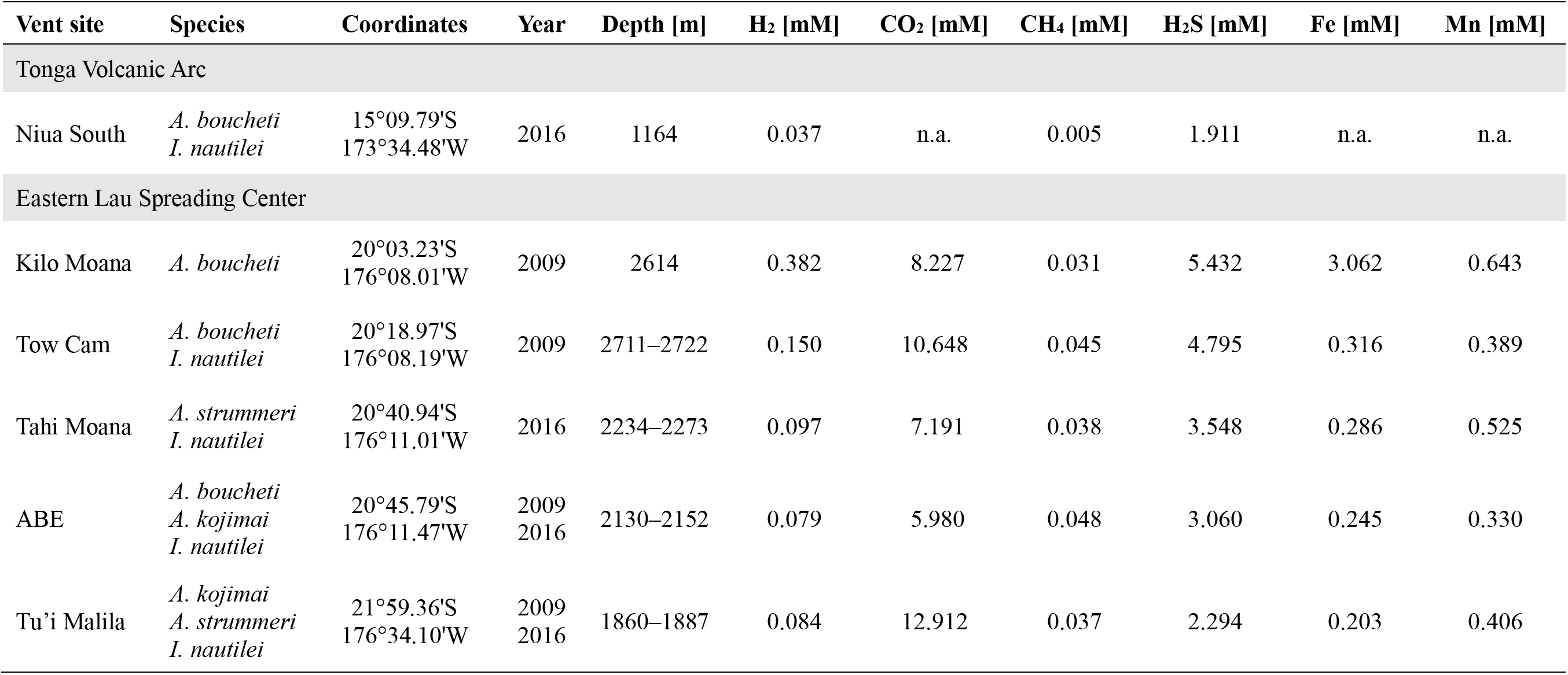
Sampling information for *Alviniconcha* and *Ifremeria* species from the Eastern Lau Spreading Center and Tonga Volcanic Arc. Chemical concentrations are mean endmember values for each vent field obtained from Beinart et al. [23], Mottl et al. [88], Flores et al. [89], and unpublished data provided by J. Seewald (Woods Hole Oceanographic Institution) and A. Diehl (MARUM). n.a. = not available.

## Results

### Host transcriptome assemblies and population genomic structure

Host transcriptome assemblies consisted of 24,176–35,654 transcripts (totaling 20.68–28.68 Mb) and were approximately 30.30–55.40% complete (Table S1). Mapping of host reads against the transcriptome references and subsequent filtering of variant sites yielded 1,655–9,185 single nucleotide polymorphisms (SNPs) per species for population genetic analyses. Irrespective of host taxon, F_ST_ and ordination analyses revealed low genetic differentiation among host populations sampled from different vent localities (Fig. 2; Table S2). With the exception of four SNP sites in *A. kojimai*, no F_ST_ outliers could be detected in any host species (Table S3). However, all species contained a number of SNPs that were moderately to highly divergent among host populations. When analyses were constrained to these SNP subsets (*Alviniconcha*: F_ST_ > 0.15; *Ifremeria*: F_ST_ > 0.10), population genetic structuring by vent site was observed in all *Alviniconcha* species, but not *Ifremeria* (Fig. S1). For both *A. kojimai* and *A. strummeri*, the respective SNP subsets further indicated genetic differences between Tui Malila populations that were sampled in different years. The effect of geography on host population genetic structuring was not significant after correction for symbiont genetic distance in partial Mantel correlations (Table S4).

**Figure 2.**
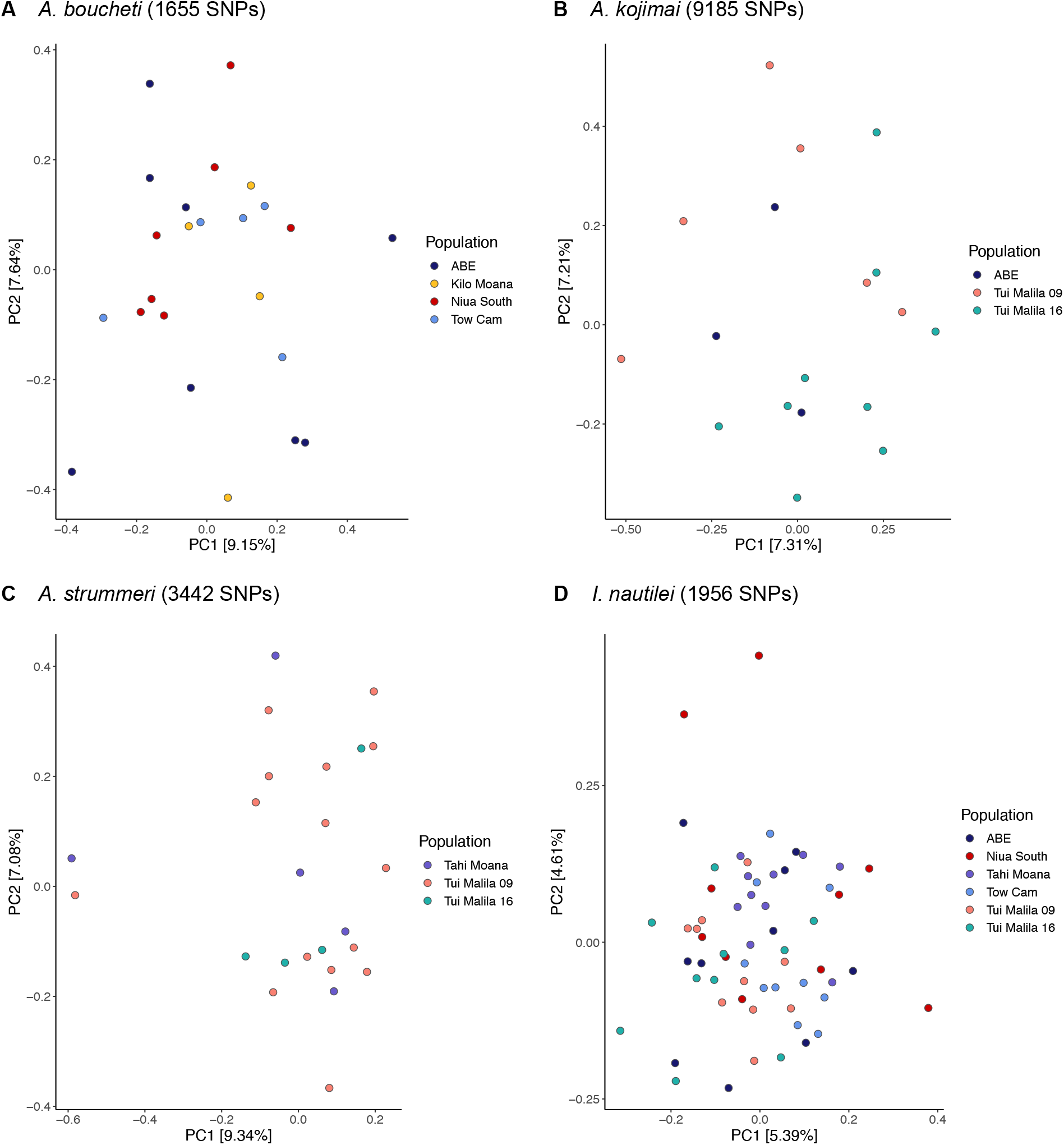
Principal component plots for *Alviniconcha* and *Ifremeria* host species based on genetic covariance matrices.

### Symbiont genome assemblies and population genomic structure

Symbiont genome assemblies varied in size from 2.10–4.86 Mb and contained 2,006–6,660 predicted protein-coding genes, with GC content ranging from 34.30–59.00% (Table S2). All assemblies were characterized by a high level of completeness (92.79–99.25%) and low amount of contamination (0.74–5.48%) (Table S5). Symbiont populations for the Epsilon, Gamma1, Ifr-SOX and Ifr-MOX phylotypes were largely structured by vent field or broader geographic region based on 239–7,057 variant sites (Fig. 3, S2; Table S6). These associations were significant even when corrected for host genetic variation based on highly differentiated SNP markers (Table S4; *p* ≦ 0.0244, *r* = 0.2665–0.8705). Genetic differentiation between symbiont populations typically increased with geographic distance between vent fields (Table S6). Although F_ST_ values for GammaLau populations of *A. strummeri* were moderate when calculated between distinct vent sites (Table S6), ordination analyses and Mantel tests provided no evidence for genetic differentiation of this symbiont phylotype across geographic locations based on 192 marker loci (Fig. 3, S2). The Gamma1 phylotype is associated with both *A. kojimai* and *A. strummeri* and we therefore investigated whether populations of this phylotype varied between host species. Ordination analyses indicated a clear clustering by host taxon that superseded the effect of geography (Fig. 4, S3), suggesting that different Gamma1 strains associate selectively with either *A. kojimai* or *A. strummeri*. By contrast, the effect of host genetics on symbiont genetic variation within species appeared to be weak. Although partial Mantel tests suggested significant associations of host genotype with strain composition or dominant symbiont type for the *A. boucheti* – Epsilon, *A. kojimai* – Gamma1, *A. strummeri* – Gamma1, and *I. nautilei* – Ifr-SOX pairs, *r* statistics were relatively low especially compared to the effect of geography, indicating limited biological relevance (Table S4; *p* ≦ 0.0364, *r* = 0.0510–0.2270).

**Figure 3.**
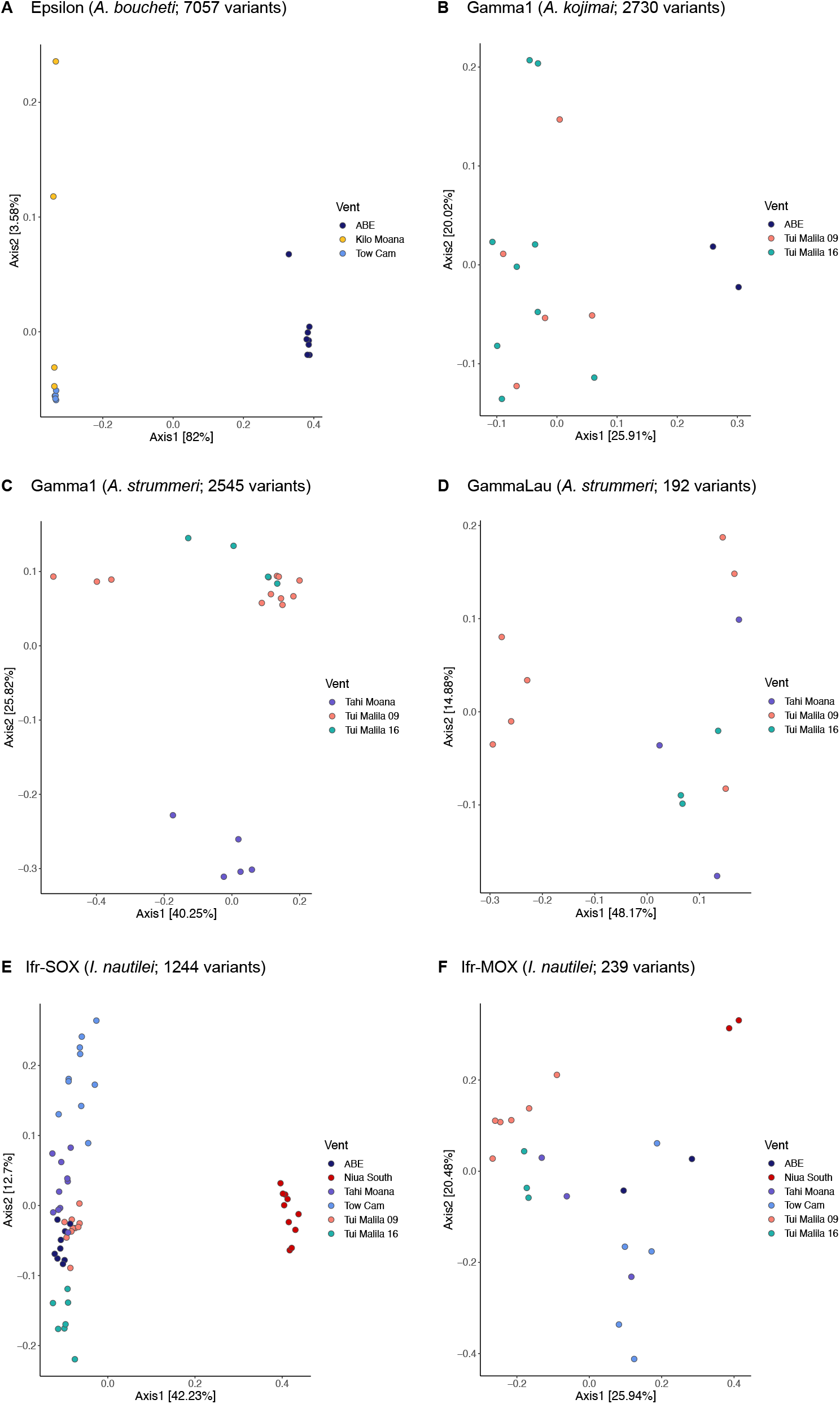
Principal coordinate plots for *Alviniconcha* and *Ifremeria* symbionts based on relative allele counts transformed into Bray-Curtis dissimilarities. Allele counts approximate the relative proportions of different symbiont strains within host individuals.

**Figure 4.**
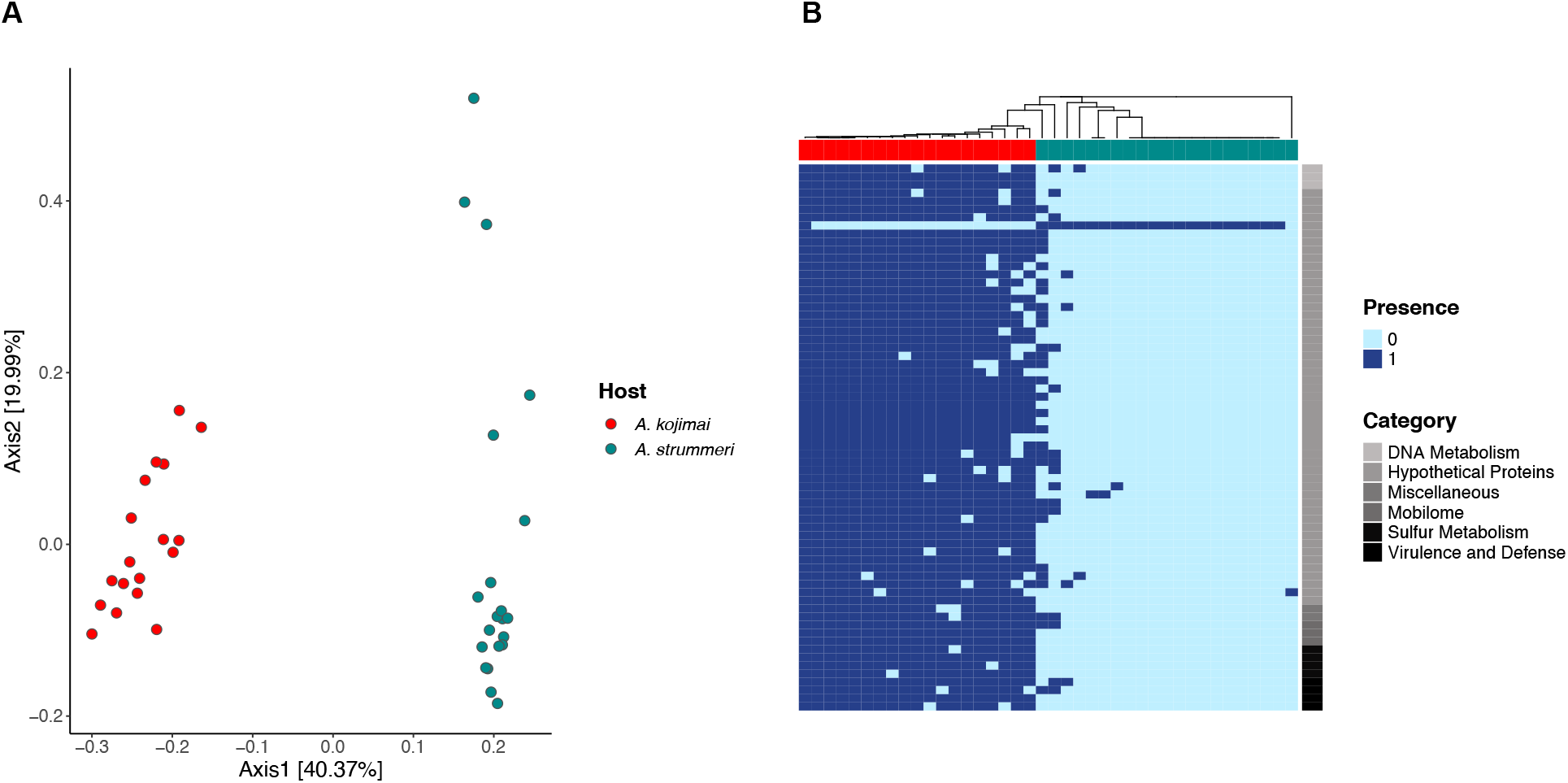
Principal coordinate plot based on Bray-Curtis dissimilarities (A) and presence/absence heatmap of differentially preserved genes (B) for the Gamma1 symbiont of *A. kojimai* and *A. strummeri*. Strains of this symbiont are clearly distinct between the two host species, even when these taxa co-occur (see Fig. S3). Segregation of symbiont populations by host species along the first ordination axis suggests that host affinity is a stronger predictor than geography for symbiont composition between species. Dark and light blue colors in the heatmap indicate presence and absence of genes, respectively. Dendrograms show similarities between samples based on their gene content profiles.

### Gene content variation between symbiont strains

We assessed variation in gene content between symbiont strains from different vent localities or broader geographic regions (Fig. 5, S4; Table S7). For the Gamma1 symbiont, we further determined gene content variation between strains from different host species given that this symbiont phylotype occurs in both *A. strummeri* and *A. kojimai* (Fig. 4, S3; Table S8). Differentially preserved or abundant genes between symbiont strains comprised about 1–5% of protein-coding regions. Independent of symbiont phylotype, the most common differences among geographic or host-specific strains concerned genes of unknown function as well as a smaller number of genes related to mobilome and anti-viral defense (Fig. 4–5, S3–4; Table S7, S8). Geographic strains of the *A. boucheti* Epsilon symbiont further differed in the presence of an ABC transporter, a GDP-L-fucose synthetase, a NAD(FAD)-utilizing hydrogenase and the DNA repair protein RecN, which were conserved in strains from Kilo Moana and Tow Cam but were absent or very lowly abundant in strains from ABE (Table S7). Within the *A. kojimai* Gamma1 phylotype, strains from ABE (but rarely Tui Malila) contained a few genes involved in DNA, protein and cell wall metabolism as well as maturation and regulation of uptake (NiFe) hydrogenases (*hyaC*, *hyaD, hoxJ*). Differences in hydrogenase-related genes were also observed in the Gamma1 symbiont of *A. strummeri*, where an operon for a hydrogen-sensing hydrogenase (*hupUV*/*hoxBC*) and genes for hydrogenase maturation and assembly proteins (*hypF*, *hyaF*, *hoxV*/*hupK*, *hypD*, *hypE*, *hoxX*) were largely missing in strains from Tui Malila (but not Tahi Moana) (Table S7). Between host species, Gamma1 strains notably differed in the presence of genes for a sulfite dehydrogenase complex (*soeABC*), which was abundant in strains specific to *A. kojimai* but not *A. strummeri* (Table S8). Within the *A. strummeri* GammaLau phylotype, strains from Tui Malila contained a broad range of metabolic genes that were absent or infrequent in strains from Tahi Moana, including genes related to macronutrient metabolism, stress response, membrane transport and several other functional processes (Table S7). The Ifr-SOX strains of *I. nautilei* differed in several genes related to DNA, nitrogen, lipid and amino acid metabolism, membrane transport, cell regulation and detoxification, which were present in strains from the ELSC but mostly missing in strains from Niua South. Within the Ifr-MOX phylotype, strains from Tahi Moana and Tui Malila differed in genes involved in sulfide respiration and oxidative phosphorylation, macronutrient metabolism, cation transport, motility, cofactor, cell wall, DNA and nucleotide metabolism as well as cell regulation and stress response (Table S7).

**Figure 5.**
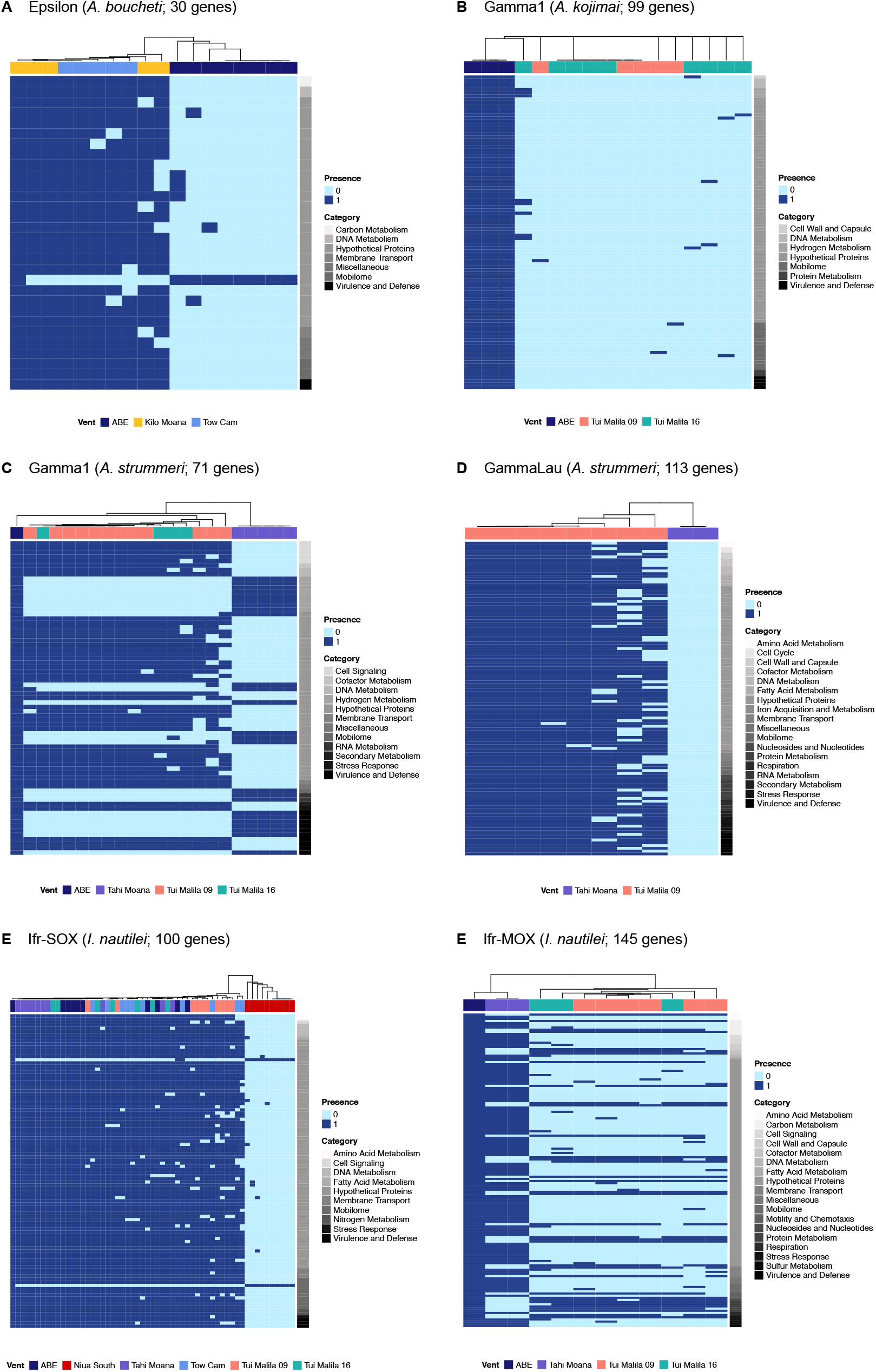
Presence/absence heatmap for differentially preserved genes.

### Adaptive variation within symbiont phylotypes

A subset of genetic variants exhibited significant associations with environmental factors in all symbiont phylotypes except for GammaLau, with constrained ordinations explaining between 5.39% and 19.76% of the variance in the respective RDA models (Fig. 6; Table S9). Candidate adaptive loci for each symbiont phylotype encompassed a variety of metabolic categories, although in many cases no particular function could be assigned (Table S9). In the *A. boucheti* Epsilon symbiont, 129 variants were significantly associated with fluid composition, while another 47 were correlated with depth. These variants were mostly located in genes related to protein, amino acid, and cell wall metabolism, followed by cofactor, DNA, carbon and nucleotide metabolism, as well as membrane transport (Table S9). Several other variants were linked to nitrogen, iron and RNA metabolism, virulence, motility, cell signaling, stress response, respiration, cell cycle, as well as hydrogen and sulfur metabolism. In the *A. kojimai* Gamma1 symbiont, 91 variants were correlated with fluid composition and 9 were linked to year. 29 of these variants were classified into functional categories. The most represented categories were mobilome, cell wall and DNA metabolism, anti-viral defense, membrane transport, cofactor metabolism and respiration. A few variants were associated with amino acid, hydrogen, protein and RNA metabolism, and cell regulation. In the *A. strummeri* Gamma1 symbiont, 95 variants showed associations with fluid composition, while 3 were correlated with year. 54 of these variants could be functionally annotated and assigned to the following metabolic categories: protein, amino acid and cell wall metabolism, membrane transport, virulence, carbon, nitrogen, DNA/RNA and nucleoside metabolism, cofactor, fatty acid and sulfur metabolism, stress response, cell signaling and mobilome (Table S9). The *I. nautilei* Ifr-SOX and Ifr-MOX symbionts contained 94 and 11 putatively adaptive variants, respectively. In both symbionts, these variants were mostly linked to depth, followed by fluid composition and year. While the majority of variants were located in genes with unknown functions, a number of loci was linked to mobile elements, cell cycle, anti-viral defense, DNA/RNA metabolism, and membrane transport. In each symbiont phylotype, virtually all candidate variants were characterized by elevated F_ST_ and gene-wide *p*N/*p*S values, further supporting their potential role in habitat adaptation (Table S9). Several other genes exhibited increased ratios of non-synonymous to synonymous substitution, although sites within these genes did not show significant associations with environment (Table S10). It is possible that sites within these genes covary with other ecological factors that could not be tested in this study or that these patterns are caused by recent slightly deleterious mutations that were not yet purged by natural selection.

**Figure 6.**
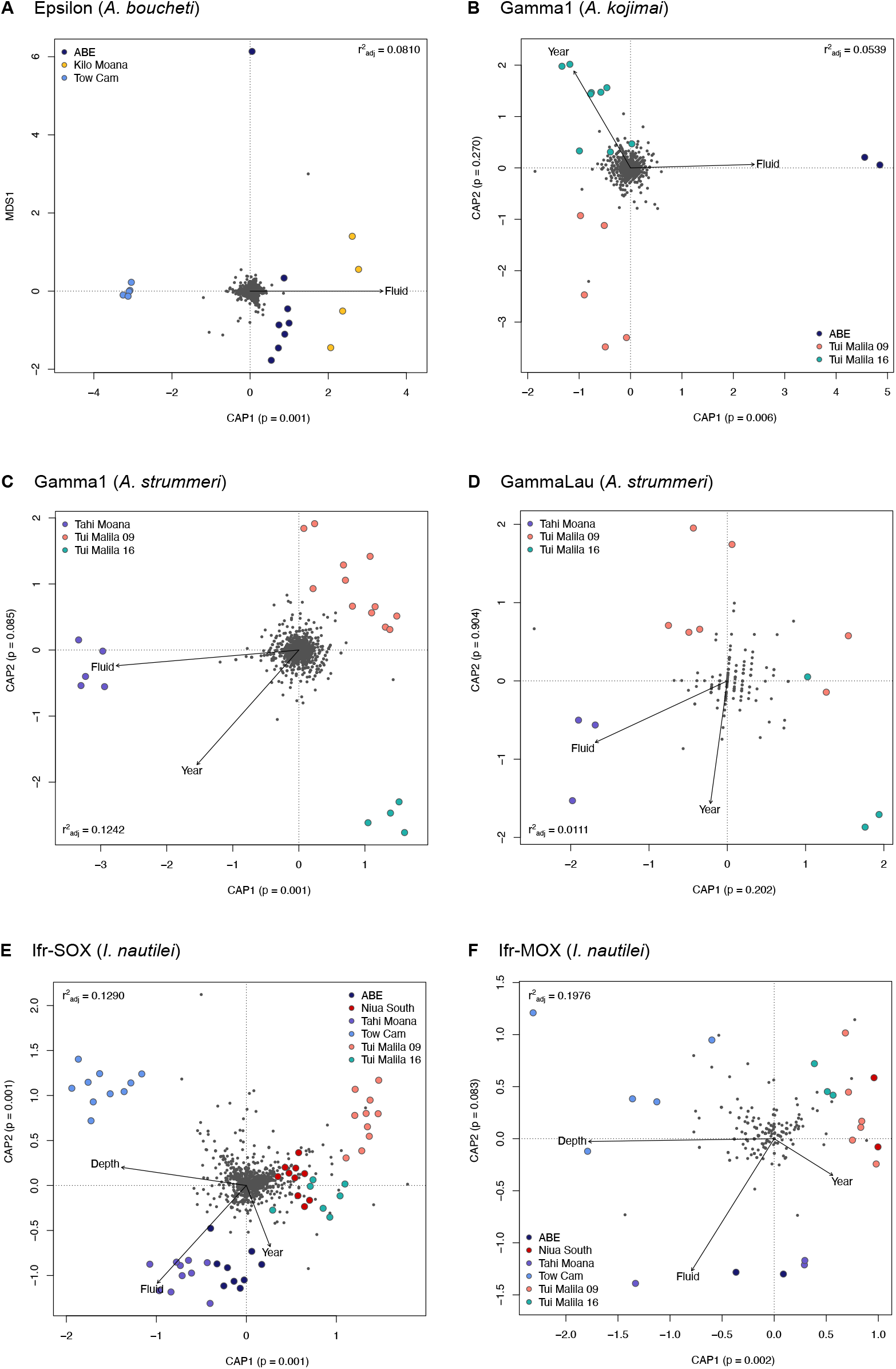
Redundancy analysis plots of the first two ordination axes for *Alviniconcha* and *Ifremeria* symbionts. Small grey dots in the center of the plot indicate genetic variants, while large colored dots represent intra-host symbiont populations sampled from different vent localities. Vectors show the environmental predictors. Redundancy analyses were conditioned by geography for all symbionts except for Gamma1 and GammaLau. Genotype-environment associations were not significant for GammaLau.

## Discussion

Strain-level variation within microbial symbionts is increasingly recognized as an important driver of host ecology and evolution (6), yet its patterns, determining factors and functional implications remain poorly investigated in many non-model symbioses. In this study, we used metagenomic analyses to assess sequence and gene content differences between chemosynthetic symbionts associated with four co-occurring species of deep-sea hydrothermal vent snails and determined the impact of host genetic, geographical and environmental factors on symbiont strain composition and variation.

Despite fidelity between hosts and symbionts at the species level (23, 25), our results indicate that specificity between host genotypes and symbiont strains in *Alviniconcha* and *Ifremeria* is weak: Host populations were not partitioned across a ~800 km gradient, whereas symbiont populations were clearly structured between vent locations or broader geographic regions. Even when we correlated symbiont and host genetic distances based on highly differentiated markers, test statistics for significant associations remained low, suggesting that host genetics has a minor impact compared to environment or geography on strain composition within both *Alviniconcha* and *Ifremeria*.

These findings qualitatively agree with observations in deep-sea mussel hybrids that appear to associate with locally available symbiont strains (19), but contrast markedly with patterns in other horizontally transmitted partnerships, such as the squid-*Vibrio* symbiosis and some legume-rhizobia associations, where hosts exhibit strong strain specificity (6). Environmental uptake of locally adapted symbiont strains provides host organisms with the opportunity to optimally exploit novel habitats, but carries the risk of unsuccessful symbiont acquisition and infection by cheaters (3). Depending on the amount of partner reliance, chances of symbiont encounter and fitness variation among symbiont strains, holobionts likely find different tradeoffs between these opposing factors. Associations with chemosynthetic bacteria are obligate for hydrothermal vent animals, but are restricted to relatively ephemeral habitats that are characterized by large temporal and spatial fluctuations in environmental conditions and associated shifts in microbial communities (29–31). Strong nutritional dependency in vent symbioses combined with the uncertainty of habitat (and thus symbiont) encounter might promote specificity towards a mutualistic symbiont phylotype, while enabling enough flexibility towards different local strains of that phylotype to maximize recruitment success of host larvae at a new vent site. By contrast, bobtail squids inoculate their environment with symbiotic bacteria and thereby ensure symbiont availability for their offspring (32), which might favor increased strain selectivity in this association. Similarly, some legume species appear to preferentially associate with certain rhizobial strains that are abundant in the native host range (33). Although the factors underlying strain specificity are not fully understood, it is possible that the relative fitness advantages provided by these locally available strains (33) coupled with the benefit of decreased cheater invasion (11) might have selected for strong partner fidelity in these symbioses.

*Alviniconcha kojimai* and *A. strummeri* represent notable exceptions to the observed patterns, as they share identical phylotypes of one gammaproteobacterial symbiont (Gamma1), but appear to take up different strains even in habitats where these host species co-occur. The phylogenetic divergence between *A. kojimai* and *A. strummeri* (~25 MYA) is more recent than that between either species and *A. boucheti* (~38 MYA) or *I. nautilei* (~113 MYA) (25). Perhaps the closer evolutionary relationship between these two species favors associations with the same symbiont phylotype, as has been suggested for *Bathymodiolus azoricus* and *B. puteoserpentis* on the Mid-Atlantic Ridge as well as *B. thermophilus* and *B. antarcticus* on the East Pacific Rise (19). However, compared to the bathymodiolin mussel system, the split between *A. kojimai* and *A. strummeri* is significantly older (>17 MYR), which could indicate that the timing of phylotype specificity evolution varies among taxonomic groups, possibly as a result of contrasting ecological or evolutionary contexts. The fact that *A. kojimai* and *A. strummeri* nevertheless associate with distinct symbiont strains in sympatry is potentially a mechanism to avoid competition for niche space. Gamma1 strains of these two species notably differed in the presence of a molybdenum-containing sulfite dehydrogenase (SoeABC), which was preserved in strains of *A. kojimai* but not *A. strummeri*. SoeABC is the predominant enzyme for sulfite oxidation in many purple sulfur bacteria (34) and is further involved in taurine and dimethylsulfoniopropionate degradation in *Roseobacter* clade bacteria (35–37). Although the physiological role of SoeABC in *A. kojimai*’s Gamma1 strain is unknown, it is possible that it contributes to partitioning of sulfur resources among co-occurring snail holobionts.

Our data support a model of horizontal symbiont transmission in both *Alviniconcha* and *Ifremeria*. While these results are expected for *Alviniconcha* which produce free-swimming planktotrophic larvae and do not invest in their young (38), they are rather surprising for *Ifremeria* which brood their offspring in a modified pouch in the female’s foot (22) and would thus have the opportunity to pseudo-vertically transmit their symbionts. It is possible that a pseudo-vertical transmission component exists in *Ifremeria*, but that maternally acquired symbionts get replaced or complemented by more competitive strains in the habitat where the snail larvae settle. Our current dataset cannot distinguish this possibility from a strict horizontal transmission mode, though future studies assessing symbiont composition in different developmental stages of *Ifremeria* would be helpful to address this hypothesis.

While the majority of genetic variants in the symbionts did not deviate from neutral expectations, a subset of loci showed evidence for natural selection, implying a role of both genetic drift and local adaptation in shaping symbiont population structure. Candidate adaptive loci spanned a surprisingly broad range of metabolic functions, many of which were probably not causally linked to the investigated environmental predictors but other correlated variables that we could not account for in this study. For example, in almost all symbiont phylotypes we observed adaptive variation in genes that were related to anti-viral defense (e.g., CRISPR-associated proteins, restriction-modification systems) or mobile elements. Although differences in depth or geochemistry might contribute to these patterns, it is more likely that they reflect exposure of symbiont strains to distinct viral assemblages that might covary with local habitat conditions, as has been suggested for vestimentiferan tubeworm symbionts (39). Polymorphisms in other genes, by contrast, are likely directly explained by variation in the analyzed environmental factors. Geographic strains of *A. boucheti*’s Epsilon symbiont, for instance, contained several variants under positive selection that were located in genes involved in hydrogenase assembly, iron transport, sulfur oxidation and respiration. Likewise, the Gamma1 strains of *A. kojimai* and *A. strummeri* showed adaptive differences in variants related to hydrogen and sulfur metabolism, respectively. Niche-specific differences in hydrogen metabolic genes were also observed in comparisons of gene content among symbiont strains. In both *A. kojimai* and *A. strummeri*, Gamma1 strains from Tui Malila lacked some subunits of uptake or hydrogen-sensing hydrogenases as well as various genes for hydrogenase maturation, synthesis and regulation. Given that H_2_ concentrations at Tui Malila can drop to 35 μM in endmember fluids and are probably lower in diffuse flow habitats (23), it is possible that hydrogen does not constitute a major energy source for Gamma1 strains from this locality. This hypothesis is in agreement with previous physiological experiments that revealed strikingly low hydrogen oxidation and associated carbon fixation rates in Gamma1 symbionts from Tui Malila (26). An alternative though mutually non-exclusive explanation for the loss of hydrogenase-related genes at least in the *A. strummeri* strain from Tui Malila could be avoidance of intra-host competition with co-occurring GammaLau strains, which often co-dominate in *A. strummeri* individuals at this vent site (23). Such functional diversity is predicted to enable symbiont coexistence in a variety of hydrothermal vent symbioses, including bathymodiolin mussels (40, 41), alvinocaridid shrimp (42) and vestimentiferan tubeworms (43).

Compared to their symbionts, host populations were markedly less structured across the same spatial scales. Although a proportion of genetic markers was differentiated across vent sites, we did not find strong evidence for local adaptation in any host species, suggesting that these patterns likely reflect random variation among localities. The limited genetic subdivision between host populations agrees with predictions from biophysical models that indicate absence of physical dispersal barriers for vent larvae within the Lau Back-Arc Basin (44). While symbiont population structure, by contrast, appeared to be at least partly driven by natural selection, it is possible that symbionts also experience stronger dispersal limitations than their hosts, as has been hypothesized in some coral-algae symbioses (18). The environmental distributions and life cycles of the free-living stages of *Alviniconcha* and *Ifremeria* symbionts are currently unknown and a better understanding of these aspects will be necessary to evaluate the relative importance of dispersal barriers on symbiont biogeography.

Overall, our findings reveal a lack of strain-level specificity in *Alviniconcha* and *Ifremeria* symbioses, which possibly reflects an evolutionary strategy to cope with the transient and dynamic nature of hydrothermal vent habitats. Strain flexibility in these associations likely contributes to the wide-spread genetic connectivity observed among host populations, which in turn could favor ecological resilience to natural but also anthropogenic environmental disturbances, a relevant consideration given the increasing human pressures on hydrothermal ecosystems worldwide (45). Our observations further support the fundamental hypothesis that horizontal transmission in marine symbioses enables host organisms to associate with locally adapted symbiont strains (3, 12–14). Though the genomic basis of local adaptation can be detected in natural populations using population genomics methods, as we did here, evaluation of the phenotypic consequences of the observed strain-level genomic trait variation will be necessary to confirm local adaptation in these symbiont strains. Future work using organism-based manipulative experiments will be helpful to compare the fitness of hydrothermal vent animals hosting site-specific symbiont strains when exposed to native and foreign conditions (46). These assessments will be critical to understand the commonality of horizontal symbiont transmission in the marine environment, given the hypothesis that local adaptation is less common in marine than terrestrial systems due to higher levels of gene flow (47).

## Materials and Methods

### Sample collection, nucleic acid extraction and sequencing

Samples of *Alviniconcha* and *Ifremeria* were collected from six vent sites (1164–2722 m) of the Lau Basin and Tonga Volcanic Arc in 2009 and 2016 using remotely operated vehicles (Fig. 1; Table 1). Upon recovery, animal samples were dissected, placed in RNALater™ Stabilization Solution (Thermo Fisher Scientific, Inc.) and frozen at –80°C until further analysis. DNA was extracted with the Quick-DNA 96 Plus extraction kit (Zymo Research, Inc.) and further purified with the MO BIO PowerClean DNA Pro Clean-Up kit (Qiagen, Inc.). High molecular weight (HMW) DNA for long-read sequencing was isolated with Qiagen Genomic-tips following manufacturer’s instructions.

### Host transcriptome sequencing and assembly

Illumina RNAseq reads for host transcriptome assemblies were obtained from sequencing experiments performed in (48) and (26). Adapter clipping, quality trimming, and removal of rRNA and symbiont reads was performed as in (26). Cleaned host reads were error corrected with Rcorrector (49) and filtered for uncorrectable and overrepresented sequences with the TranscriptomeAssemblyTools package (https://github.com/harvardinformatics/TranscriptomeAssemblyTools). Host transcriptome co-assemblies for each *Alviniconcha* and *Ifremeria* species were performed with Trinity (50) using the PasaFly algorithm. For each *Alviniconcha* species, additional transcripts were reconstructed from 454 reads obtained from (51). Assembled contigs for each species were clustered with Cd-Hit-Est (52) at a 95% identity threshold to reduce transcript redundancies. Open reading frames (ORFs) were predicted with TransDecoder (https://github.com/TransDecoder/TransDecoder) considering homologies to known proteins (UniRef90) and protein domains (Pfam) as ORF retention criteria. Transcripts that did not contain any ORF or had a non-eukaryotic origin based on taxonomy classifications with BlobTools (53) were removed from the assembly. Final transcriptome assemblies were evaluated for quality and completeness with Busco (54).

### Symbiont genome sequencing and re-assembly

Illumina reads for the *A. boucheti*, *A. kojimai* and *Ifremeria* holobionts were obtained from previous sequencing runs performed in (27). Raw reads were trimmed with Trimmomatic (55) and filtered for sequence contaminants through mapping against the human (GRCh38) and PhiX reference genomes. Decontaminated reads were grouped into symbiont and host reads by assessing best matches against draft symbiont genomes (27) with BBsplit (https://sourceforge.net/projects/bbmap/). To improve contiguity of the genome assemblies we conducted 3–4 Nanopore sequencing runs for each symbiont phylotype on a MinION device (Oxford Nanopore Technologies) using the SQK-LSK109 ligation kit after HMW DNA enrichment with the Circulomics Short Read Eliminator kit. Basecalling of the Nanopore reads was done with Albacore (Oxford Nanopore Technologies) and adapters were clipped with PoreChop (https://github.com/rrwick/Porechop). Hybrid assemblies of Illumina and Nanopore reads were constructed for each symbiont genome with SPAdes (Gamma1) (56) or metaSPAdes (all others) (57) choosing k-mers between 21 and 91 in 10 step increments. Raw assemblies for the Epsilon, Gamma1 and Ifr-SOX symbionts were binned with GBTools (58) and incrementally gap filled, corrected and scaffolded with LR_GapCloser (59), ra2.py (60), and SLR (61), respectively, following recommendations by (62) The Ifr-MOX symbiont assembly was automatically binned with MetaBAT2 (63) for contigs ≧ 1500 bp. Shorter contigs (≧ 500 bp) were binned with MaxBin (64). Scaffolds < 200 bp were excluded from all assemblies. Final assemblies were polished with Pilon (65), functionally annotated with Rast-Tk (66) and quality-checked with CheckM (67) and Quast (68).

### Population-level metagenomic sequencing and variant identification in hosts and symbionts

192 barcoded high-throughput DNA sequencing libraries were prepared with a Tn5 transposase-based protocol after (69) at the University of California Santa Cruz and then sent for 150 bp paired-end sequencing on a NovaSeq 6000 instrument at the University of California Davis. However, due to low read allocation 66 of the libraries were excluded from further analysis. Raw sequence reads were trimmed, filtered and sorted by host species and symbiont phylotype as described above. Optical duplicates were removed with Picard’s MarkDuplicates tool (https://github.com/broadinstitute/picard). To resolve common alignment errors and improve base call accuracy, we locally realigned reads around indels and recalibrated base quality scores with LoFreq (70).

Host population genomic variation was assessed in Angsd (71) by inferring genotype likelihoods based on Hardy-Weinberg equilibrium considering individual inbreeding coefficients. To increase accuracy of the analyses, variant sites with mapping qualities < 30 (minMapQ = 30), base qualities < 20 (minQ = 20), and minimum minor allele frequencies < 0.01 (minMaf = 0.01) were excluded. We further filtered sites based on strand bias (sb_pval = 0.05), heterozygote bias (hetbias_pval = 0.05) and probability of being variable (SNP_pval = 1e-6). In addition, we removed spurious and improperly paired reads, adjusted mapping qualities for excessive mismatches (C = 50) and computed per-base alignment qualities (BAQ = 1) to disregard variants close to indel regions. Putative paralogous variants were excluded by discarding reads with multiple mappings and by considering only sites that had a maximum depth of 40–80. Genetic distances between individuals were inferred by calculating pairwise covariance matrices.

Symbiont population genomic variation was determined with FreeBayes (72) using input parameters adjusted for the analysis of metagenomic data (-F 0.01 -C 1 -p 1 --pooled-continuous --haplotype-length 0 --report-monomorphic). Variant calls were restricted to sites with a minimum base quality of 20, a minimum mapping quality of 30, and a minimum coverage of 10. To eliminate bias in variant identification and other downstream analyses due to uneven read depth between samples, we normalized all samples to the lowest amount of coverage found in a sample for a particular symbiont phylotype (> 10X coverage). Variants were further filtered based on strand bias (SRP > 5 && SAP > 5 && EPP > 5), proximity to indels (5 bp) and maximum mean depth with BcfTools (73) and VcfTools (74). In addition, sites and individuals with more than 25% missing data were excluded from the analysis. Allele counts (= symbiont strain abundances) and consensus haplotypes (= dominant symbiont strains) were extracted with GATK’s VariantsToTable tool (75).

### Genomic structure and differentiation

We performed ordination analyses with the Ape and Stats packages in R (76, 77) to assess genetic variation between symbiont and host populations. Host genetic structure was determined through principal component analyses based on genetic covariance matrices, while symbiont genetic structure was inferred through principal coordinate analyses based on both consensus haplotype and allele count data transformed into Euclidean distances and Bray-Curtis dissimilarities, respectively. For each sample, absolute allele counts were normalized to relative counts prior to analysis. Negative eigenvalues were corrected using the method by Cailliez (78) and final ordination plots were produced with GGplot2 (79). F_ST_ values between host and symbiont populations were calculated in Angsd and SciKit-Allel (https://github.com/cggh/scikit-allel), respectively, following the procedure in (80). For the host species, potential outlier loci under selective pressures were inferred with OutFlank (81) based on neutral F_ST_ distributions that were obtained from quasi-independent SNP subsets determined with Plink (82).

### Assessment of gene content variation

We used PanPhlAn (83) to investigate potential differences in gene composition between symbiont strains from different vent localities. Downsampled symbiont reads for each species were mapped against the corresponding symbiont reference genome using custom Rast-Tk annotations for functional categorizations. Samples were profiled for gene presence/absence based on the following parameter thresholds: --min_coverage 1 --left_max 1.70 --right_min 0.30. Gene content variation between symbiont strains was assessed by identifying genes that were present/absent in 90% of samples from one or the other geographic region. To account for gene content variation between different strains within hosts we further quantified gene abundances with Salmon (84) using the following parameters: -l IU --meta --rangeFactorizationBins 4 -- numBootstraps 1000 --seqBias --gcBias -s -u. Abundance values were normalized with the Trimmed Mean of M method using Trinity’s *abundance_estimates_to_matrix.pl* script (50, 85). Gene presence/absence and differential abundance heatmaps were produced with the ComplexHeatmap package in R (86).

### Assessment of symbiont variation based on environment and host genetics

We conducted partial Mantel tests with the Vegan package in R (87) to assess relationships between host and symbiont genetic distances and geography based on Spearman rank correlations. As no population genetic structure could be observed in any host species based on the full SNP datasets, we estimated host covariance matrices using only moderately to highly differentiated SNP subsets for this analysis (*Alviniconcha*: F_ST_ > 0.15; *Ifremeria*: F_ST_ > 0.10). Geographic distances were determined by calculating the geodesics between vent sites with the GeoSphere package (88). We used redundancy analyses (RDA) following the approach in (89) to evaluate the influence of hydrothermal fluid composition and depth on the genetic structure of each symbiont phylotype. We further included the effect of sampling year, which encompasses changes in hydrothermal circulation within the ELSC as indicated by the cessation in fluid flow at the Kilo Moana vent field between 2009 and 2016. Endmember concentrations for geochemical compounds were obtained from the literature (23, 90, 91) or unpublished data provided by A. Diehl and J. Seewald (Table 1). Due to multi-collinearity among the chemical species, we used the first eigenvector from principal coordinate analyses (corresponding to the eigenvalue with the largest explanatory power) as composite value. For each symbiont phylotype, we further assessed the strength of correlation with other predictors to exclude variables that were highly collinear. Unless geographic location was strongly linked to environmental predictors, we used latitude as a conditioning factor in the analyses to correct for isolation by distance. Based on these collinearity evaluations, we tested the effect of all factors on the two *Ifremeria* symbionts, the effect of depth and fluid composition on the *A. boucheti* symbiont and the effect of fluid composition and year (uncorrected for geography) on the *A. kojimai* and *A. strummeri* symbionts. SNPs were considered candidates for local adaptation if their loadings on significant constrained RDA axes deviated more than 2.5 standard deviations from the mean of the distribution. For each symbiont phylotype we further calculated the ratio of non-synonymous to synonymous polymorphisms (*p*N/*p*S) with SNPeff (92) to infer candidate genes evolving under natural selection.

### Data Accessibility

Raw Nanopore reads, host transcriptomes and symbiont genomes have been deposited in the National Center for Biotechnology Information under BioProjects PRJNA523619, PRJNA526236 and PRJNA741492. Annotations for new symbiont genomes are available on the Rast webserver (https://rast.nmpdr.org/rast.cgi) through the guest access (login: guest, password: guest) under job IDs 6666666.685764 (Epsilon), 6666666.685765 (Ifr-SOX), 6666666.686795 (Gamma1), and 6666666.687466 (Ifr-MOX). Scripts for bioinformatic analyses are available on GitHub under https://github.com/cbreusing/Provannid_host-symbiont_popgen.

## Acknowledgements

We thank the captains, crews and ROV pilots of the R/V *Thomas G. Thompson* (ROV *Jason II*) and R/V *Falkor* (ROV *Ropos*) for supporting the sample collections that made this study possible. We thank Peter Girguis for his contribution of the 2009 samples to this project, Alexander Diehl and Jeff Seewald for providing geochemical data, Michelle Hauer and Erin Frates for their assistance with sample preparation and the National Science Foundation EPSCoR Cooperative Agreement OIA-#1655221 for providing access to Brown University’s high-performance computing cluster where the bioinformatic analyses were performed. We further thank the technical staff at the UC Davis Genome Center for sequencing our Illumina metagenomic libraries. This work was supported by the National Science Foundation (grant numbers OCE-1536331, 1819530 and 1736932 to R.A.B.) and the National Institute of Health (grant numbers 5K99GM135583-02 to S.L.R. and 5R35GM128932-03 to R.C.D.).

