## Supplementary Information for "Host-symbiont population genomics provide insights into partner fidelity, transmission mode and habitat adaptation in deep-sea hydrothermal vent snails"

**This PDF file includes:**

**Figures S1 to S4**
**Legends for Tables S1 to S10**

**Other supplementary materials for this manuscript include the following:**

**Tables S1 to S10**

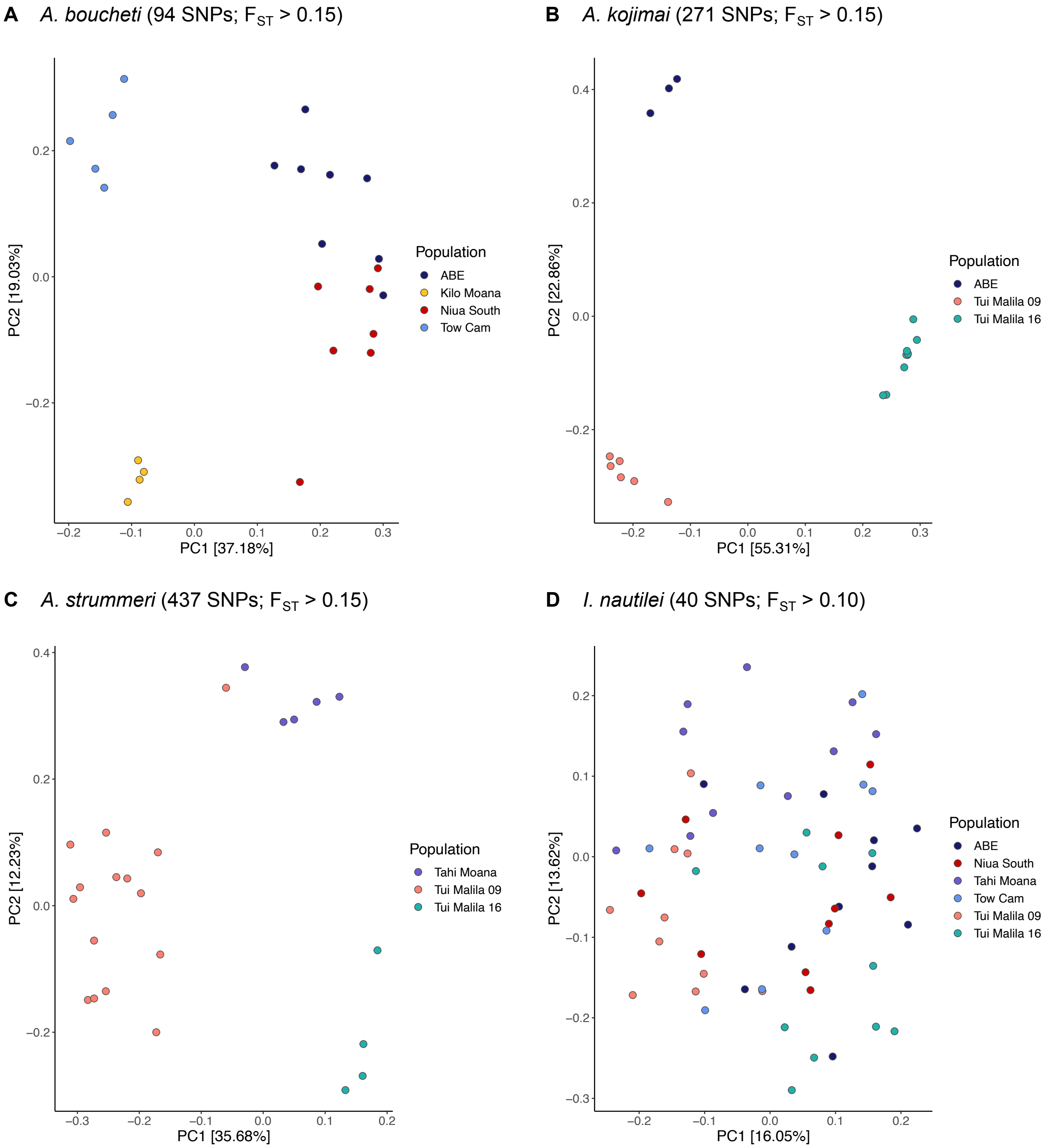


**Fig. S1** Principal component plots for *Alviniconcha* and *Ifremeria* host species based on genetic covariance matrices calculated from moderately to highly differentiated SNPs.

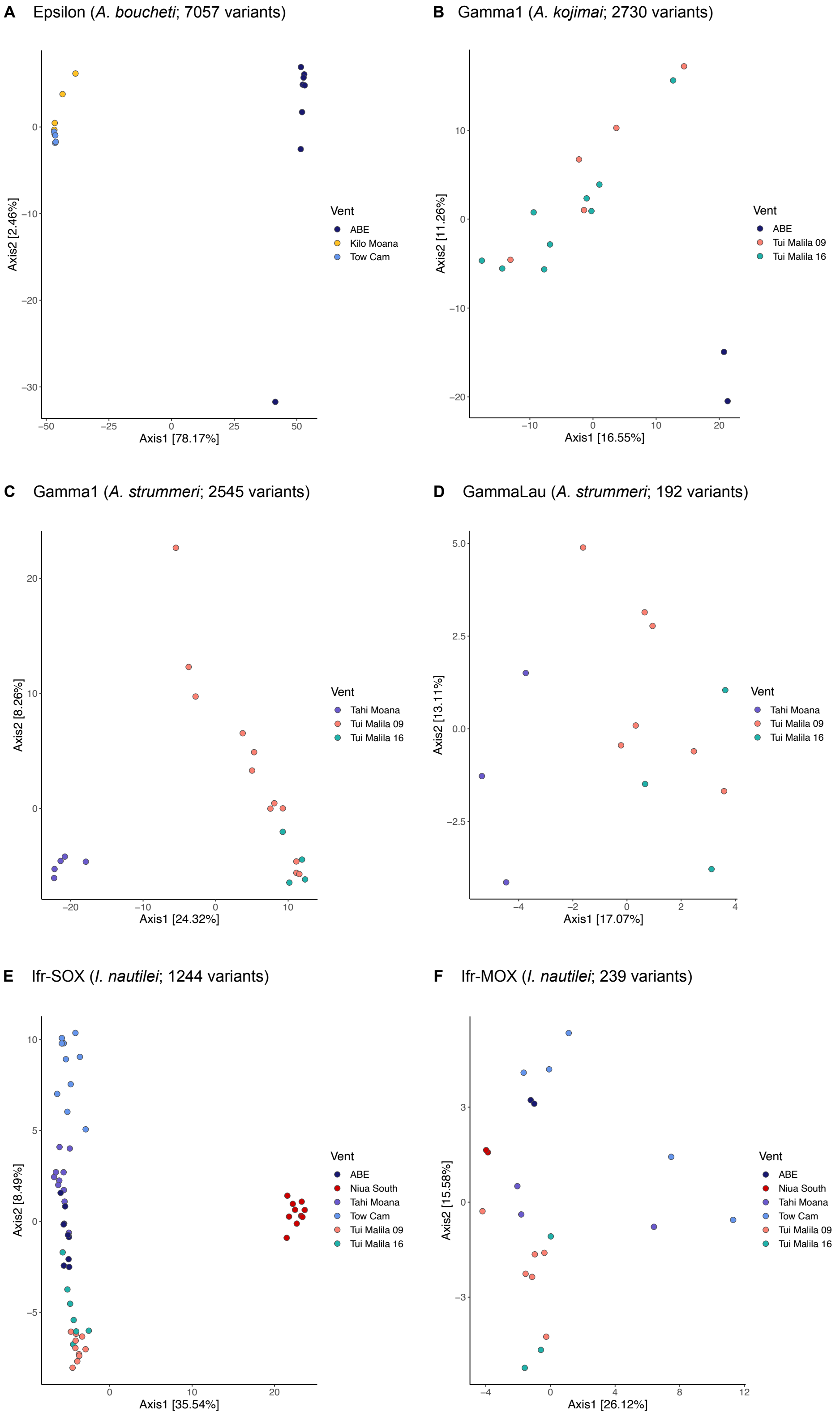


**Fig. S2** Principal coordinate plots for *Alviniconcha* and *Ifremeria* symbionts based on consensus haplotypes transformed into Euclidean distances. Consensus haplotypes represent the dominant symbiont strain within host individuals.

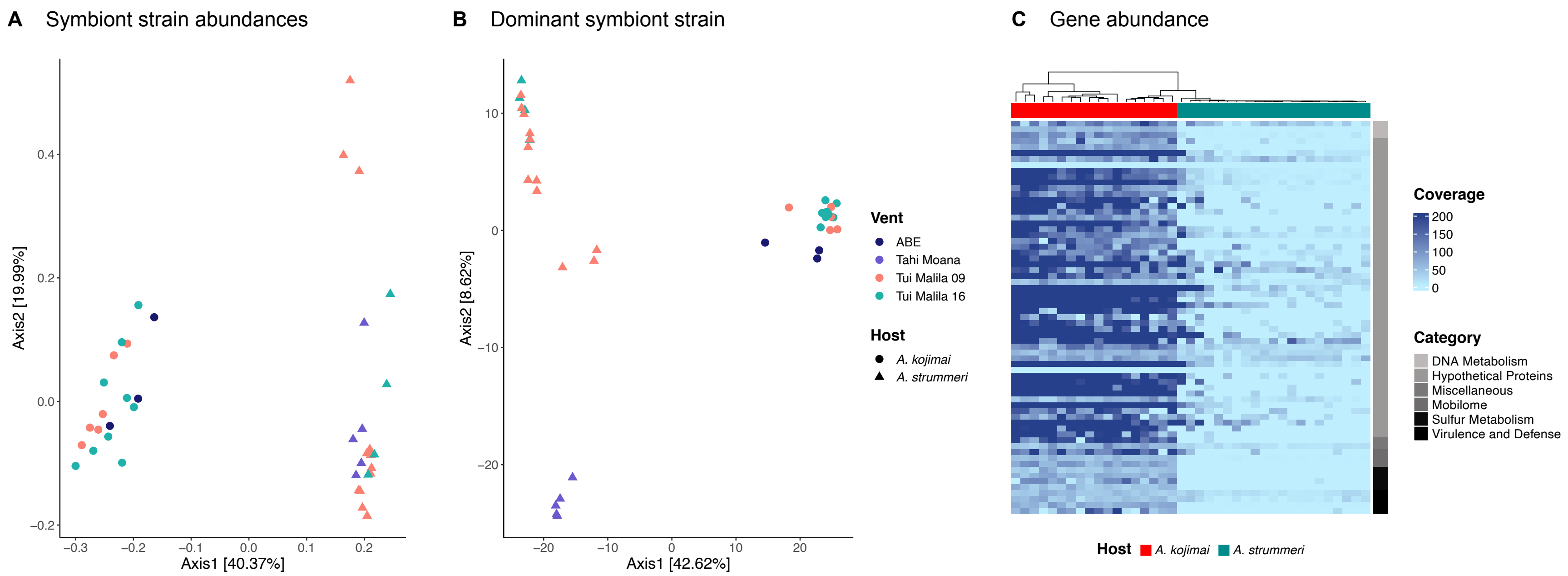


**Fig. S3** Principal coordinate plots and gene coverage heatmaps for the Gamma1 symbiont of *A. kojimai* and *A. strummeri*. (A) Principal coordinate analyses based on relative allele counts (symbiont strain abundances). (B) Principal coordinate analyses based on consensus haplotypes (dominant symbiont strain). (C) Gene abundance heatmap for differentially preserved genes.

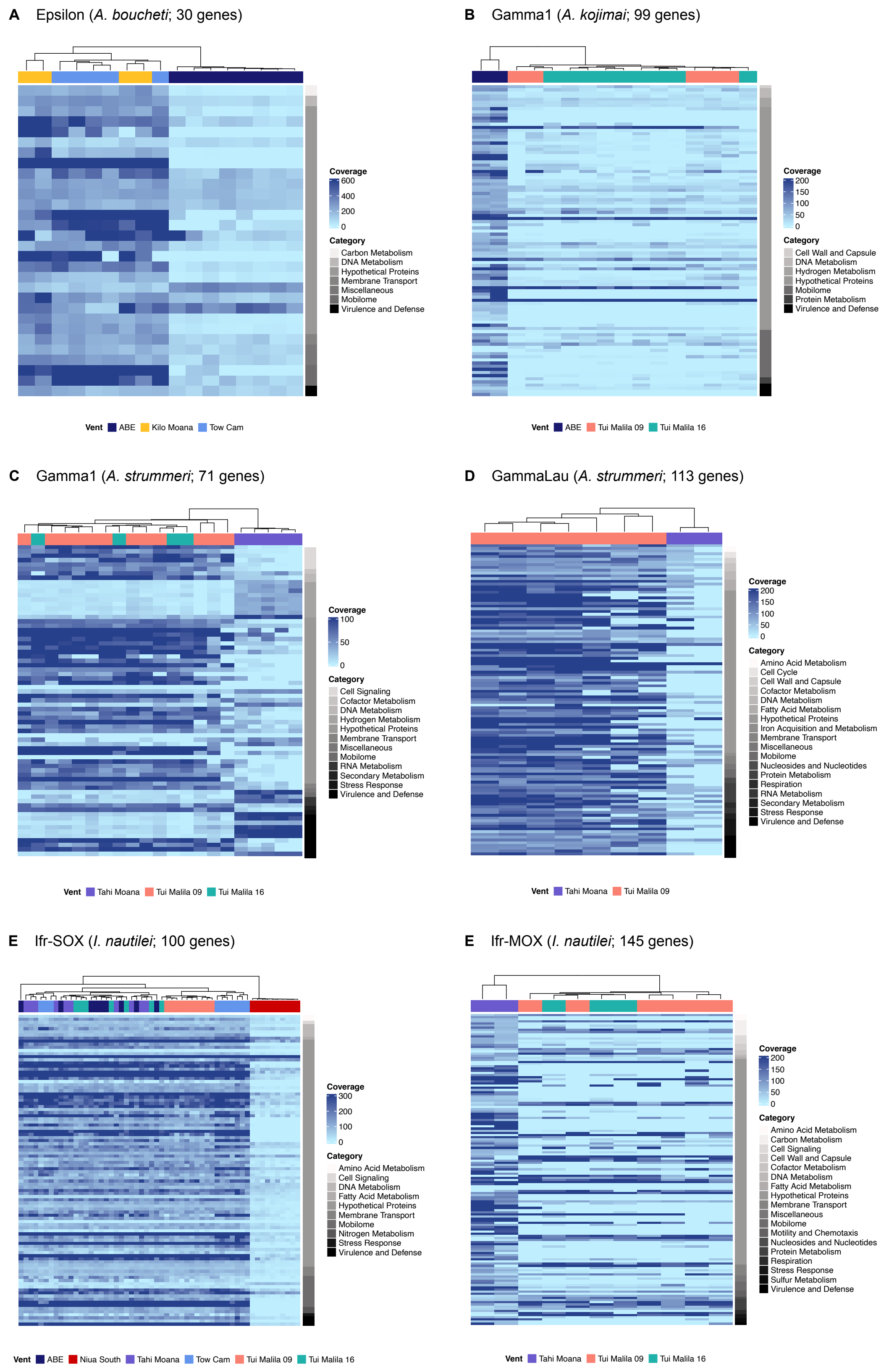


**Fig. S4** Gene abundance heatmap for differentially preserved genes in *Alviniconcha* and *Ifremeria* symbionts.

**Table S1** General statistics for host transcriptomes assembled in this study. Statistics are based on CDS regions.

**Table S2** Pairwise $F_{ST}$ values for *Alviniconcha* and *Ifremeria* populations from the Lau Basin.

**Table S3** $F_{ST}$ outlier tests and functional annotation of polymorphic transcripts in *Alviniconcha* and *Ifremeria* host species.

**Table S4** Partial Mantel tests between symbiont, host and geographic distances. Significant values are indicated in bold.

**Table S5** General statistics for symbiont genome assemblies.

**Table S6** Pairwise FST values for *Alviniconcha* and *Ifremeria* symbiont populations between vent localities.

**Table S7** Gene content variation among geographic symbiont strains.

**Table S8** Gene content variation among Gamma1 strains of *A. kojimai* and *A. strummeri*.

**Table S9** Summary of variants correlated with environmental factors.

**Table S10** *p*N/*p*S and $F_{ST}$ values for symbiont genes.
